# Dietary protein restriction in rats leads to a rapid within-session preference for protein

**DOI:** 10.1101/2025.01.24.634712

**Authors:** Giulia Chiacchierini, Fabien Naneix, John Apergis-Schoute, James E McCutcheon

## Abstract

Evolution has provided species with adaptive behavioural mechanisms that guarantee tight dietary protein regulation. However, an unsolved question is whether the well-established preference for protein-containing food manifested during states of protein restriction is innate or learned. Here, we tackled this problem by maintaining male rats on either a low-protein diet (5% protein, protein-restricted) or a control diet (22% protein, non-restricted) for 9-12 days, and then offered them two novel foods, a protein-containing solution (4% casein) and a carbohydrate-containing solution (4% maltodextrin) during a daily 60-minute free-choice test, repeated for 5 consecutive days. We assessed both the total and cumulative intake of each solution throughout each test, as well as the microstructure of licking behaviour as an index of the solutions’ palatability. In a second experiment, we exposed a different cohort of rats, before any behavioural test, to the same protein source (i.e., casein) that they would encounter during the free-choice tests, to assess whether familiarity with casein would drive subsequent casein intake even in non-restricted rats. We found that dietary protein restriction leads to a rapid preference (within 16 minutes of first exposure) for a casein-rich solution, and this preference is persistent over subsequent exposures. Increased palatability of protein during initial exposure correlated with protein preference in the restricted rats. Moreover, familiarity with casein did not lead to protein preference in non-restricted rats. This study demonstrates that, when in need of protein, protein preference is a rapid adaptation that requires minimal experience of protein.

**Highlights:**

- Protein-restricted rats show a rapid preference for protein over carbohydrate
- The preference emerges within 5 minutes of exposure to solutions
- Increased palatability of casein is partially related to preference
- Exposure to casein without protein restriction does not lead to preference

## 1. Introduction

Many animals have been shown to actively shift their food preference according to nutrient-specific needs, an essential ability for homeostasis and survival [1,2]. In particular, the need for dietary protein strongly influences food choice in many different species, from insects to humans [3,4]. Rodents, for example, increase their protein intake during periods of growth [5,6], pregnancy and lactation [7,8], in response to growth hormone [9], and as a consequence of dietary protein restriction [10–14]. Recent studies have begun to shed light on the neural and cognitive processes associated with this need-driven protein appetite but much is still unknown [15–19].

A key remaining question is whether the adaptive response to protein deficiency is innate or learning-dependant [20]. In favour of an unlearned protein appetite, it has been reported that protein-depleted rats exhibited protein preference from the first minute of their first exposure to protein-rich food [21], driven by olfactory cues [22]. A similarly fast process is observed with sodium appetite [23,24] and is associated with motivational and perceptual changes, such as increased palatability of sodium [25]. In contrast, other studies reported that protein-restricted rats needed several days to adapt their food intake towards a high-protein diet [26,27], a process requiring learning through post-ingestive feedback [28]. More recent work examined a 5-hour time frame of exposure to novel protein and carbohydrate solutions, showing that protein-restricted mice significantly increased intake of protein after 150 minutes [12]. Thus, how quickly animals select protein-rich foods when in need remains an open question as are the underlying mechanisms that would support this process.

We recently reported that protein-restricted rats showed a preference for a protein-containing solution, relative to carbohydrate [10,13,15], that was associated with increased palatability of the protein solution [10,13,29], as assessed by analysis of licking microstructure [30]. Moreover, we and others have shown that protein-restricted rats and mice also show increased motivation to obtain protein-rich food [16,18]. However, both protein preference and motivation were tested after some form of conditioning (Pavlovian flavour conditioning [10,13,29] or instrumental conditioning [16]), so we could not conclude whether the drive for protein was intrinsic or acquired.

Here, to better understand whether protein preference is innate or learned, we gave protein-restricted and control non-restricted rats a choice between a novel protein-containing solution (casein) and a novel carbohydrate solution (maltodextrin), without any prior experience of the solutions, and assessed intake and timing of consumption (Experiment 1). Further, we determined the palatability of each solution at first exposure and during subsequent exposures. Then, we assessed whether familiarity with protein-containing food (i.e., casein) would increase subsequent consumption of a casein-rich solution (Experiment 2) even in non-restricted rats.

We found that rats that were protein-restricted showed a rapid preference for the novel protein solution (within 5 minutes; Experiment 1), and that this was not seen in non-restricted rats even those that had previous experience with casein-containing food (Experiment 2).

## 2. Materials and Methods

### 2.1 Animals

Male Sprague-Dawley rats (Experiment 1, n = 28; Experiment 2, n = 16) were used for the experiments (Charles River; >275 g at the beginning of the experiment). Rats were housed 2 per cage, in standard conditions as previously described (IVCs, 21 °C ± 2 °C, 40-50% humidity, 12 h:12 h light/dark cycle (lights on at 07:00) [10,15,16]. Water and chow were available ad libitum except when rats were being tested in operant chambers. All procedures were conducted under the Animals (Scientific Procedures) Act (1986), under the appropriate license authority (Project License: 70/8069).

### 2.2 Diet manipulations

All rats were initially maintained on standard laboratory chow diet (EURodent Diet 5LF5, LabDiet; 3.4 kcal/g) containing 22% protein. Half of the rats were randomly assigned to a state of protein restriction (PR), by switching the standard chow to a modified AIN-93G diet designed to be low in protein (4% protein from casein; #D15100602, Research Diets; 4.0 kcal/g), as previously described [10]. The rest of the rats were kept under a standard laboratory chow diet (non-restricted rats, NR). Food intake data are shown in Fig. S2.

In Experiment 2, NR rats were provided with both ad libitum chow, and the following amounts of the casein-containing diet (#D15100602; 30 g per cage on the first day of diet manipulation, 20 g on day 2-4, 10 g from day 5 until the end of the experiment). This allowed NR rats to familiarise themselves with casein-containing food before undergoing the preference test sessions, but without inducing a state of protein restriction. All behavioural experiments started 9 to 12 days following protein restriction.

### 2.3 Behavioural experiments

All behavioural experiments took place within standard operant chambers (Med Associates, St. Albans City, VT) equipped with house light, two bottles and contact lickometers as previously described [10,15]. Spouts were recessed from the wall of the chamber to prevent fluid bridges forming during bursts of licks and ensure that individual licks were captured with high fidelity (Fig. S1). The onset and offset of individual licks were recorded on a computer running Med-PC IV Software Suite (Med Associates). Bottles were weighed at the beginning and end of each session and a high correlation was seen between total licks and change in bottle weight so licks were used as a proxy for intake (Fig. S1). At the end of each session All sessions lasted for one hour and took place at the start of the light period between 08:00-12:00. For the first three days, rats were trained with both bottles containing 0.2% sodium saccharin, to familiarise them with the apparatus. Following this, rats underwent one preference test session per day, for five consecutive days. During preference tests, one bottle was filled with a protein-containing solution (4% casein, 0.21% methionine, 0.2% sodium saccharin, 0.05% flavoured Kool-Aid), and the other bottle with an isocaloric carbohydrate-containing solution (4% maltodextrin, 0.2% sodium saccharin, 0.05% flavoured Kool-Aid), as previously described [10,15]. Bottle positions (left, right) and flavours (cherry, grape) associated with each macronutrient were counter-balanced between rats. All testing was conducted at the same time of day (09:00 – 13:00).

### 2.4 Microstructural analysis of solutions intake

Lick timestamps were extracted from data files and analysed in Python using custom made scripts that measured number of licks for each solution, interlick intervals, number of clusters and cluster size. Specifically, licks were divided into clusters according to interlick intervals [29,30]. Criterion for ending a cluster was an interlick interval of >500 ms.

### 2.4 Statistical analysis and data availability

Lick data for training sessions, preference test sessions, and measures of palatability (number of clusters and mean cluster size) were analysed using two-way repeated measures ANOVA, with Diet group (non-restricted vs. protein-restricted) as between-subjects factor and solution (casein vs. maltodextrin) as within-subjects factor. Protein preference was calculated as licks for casein divided by total licks and analysed using one sample t-tests vs. no preference (0.5). Protein preference of non-restricted vs. protein-restricted rats was compared using unpaired t-tests.

Cumulative lick data were analysed using linear regression to assess whether the slopes of casein and maltodextrin differed within each group. To determine at what timepoint casein preference emerged, cumulative lick data were placed into 5-minute bins and casein preference was calculated. The preference for each bin was then compared against 0.5 using one-sample t-test with the Benjamini, Krieger and Yekutieli correction method to control for False Discovery Rate (FDR) [31].

For longitudinal assessment of protein preference, two-way repeated measures ANOVA was used with Diet as between-subjects factor and Day as within-subjects factor. For change in palatability within a session and longitudinal assessment of palatability, three-way mixed ANOVA was used, with Diet as between-subject factor, and Solution and Epoch as within-subject factor. Follow-up two way ANOVAs were used where appropriate.

Casein preference and percentage change in palatability between early stage of consumption vs. late stage of each solution were correlated using Pearson correlation coefficients.

Significant effects and interactions were followed with Sidak’s multiple comparisons test. For all analyses, α was set to 0.05 and all tests were two-tailed.

In Experiment 2, the sample size was determined *a priori* using the Variable-Criteria Sequential Stopping Rule (SSR) [32,33]. To test the hypothesis that would be a difference in casein preference in protein-restricted rats but not in non-restricted rats, we elected to use one-sample t-tests on casein preference, with 0.5 as theoretical mean and α = 0.05. With an effect size of 0.84 (based on Experiment 1) and a conventional power value of 0.80, the table of stopping rules suggested a sample size of minimum 8 and maximum 24 rats, and the lower and upper bounds of the stopping criteria, 0.030 and 0.200.

Data were processed, analysed, and plotted with Python using the following packages: Pandas [34], Matplotlib [35], Pingouin [36], and Seaborn [37]. Data are available at http://doi.org/10.5281/zenodo.14334920 and custom code is available at http://www.github.com/mccutcheonlab/immediate-protein-pref.

## 3. Results

### 3.1 Protein-restricted rats rapidly show a preference for protein-containing solutions

First, we examined whether protein-restricted rats would exhibit an immediate protein preference the first time they had access to protein and carbohydrate containing solutions (casein or maltodextrin) (Fig. 1). We found that when total number of licks across the whole 1-h session was considered, protein-restricted rats drank more casein than maltodextrin, while non-restricted rats did not show any difference between solutions (Fig. 1A). Analysis of total number of licks with two-way repeated measures ANOVA revealed a Diet x Solution interaction (F(1, 26) = 5.74, p = 0.024), a main effect of Solution (F(1, 26) = 5.67, p = 0.025) and no main effect of Diet (F(1, 26) = 2.01, p = 0.168). Post-hoc Sidak’s test confirmed that protein-restricted rats (p = 0.029), but not NR rats (p = 1.000), licked more for casein than maltodextrin. Accordingly, protein-restricted rats displayed a significant preference for the casein solution over maltodextrin (one sample t-test vs. 0.5, t(13) = 3.01, p = 0.010), while non-restricted rats showed no preference (t(13) = 0.09, p = 0.934). In addition, when change in bottle weight was used as a measure of intake instead of number of licks, the results were the same (Fig. S1).

**Figure 1.**
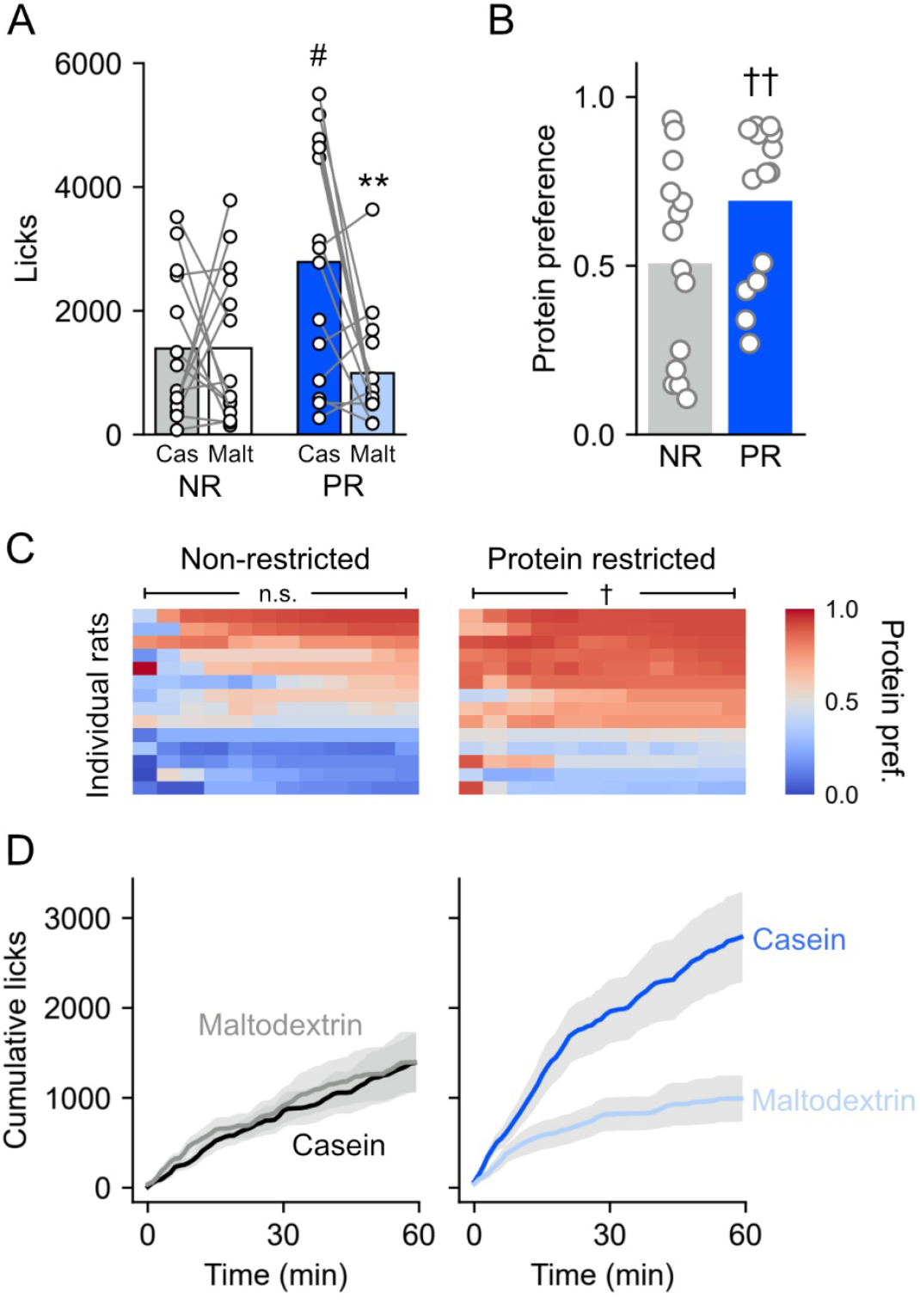
Protein-restricted rats show preference for protein over carbohydrate and the preference arises rapidly in the test. (A) Licks for casein and maltodextrin during preference test session where rats were given simultaneous access to casein and maltodextrin solutions. Protein-restricted rats increase intake of casein compared to maltodextrin and compared to non-restricted rats. Bars are mean and circles are individual rats. #, p < 0.05 vs. NR. **, p < 0.01 vs. casein (two-way repeated measures ANOVA followed by Sidak’s post hoc tests). (B) Casein preference calculated as casein licks divided by total licks. Protein-restricted (PR) rats show casein preference whereas nonrestricted (NR) rats do not. Bars are mean and circles are individual rats, †† p < 0.01 vs. 0.5 (one sample t-test). (C) Heatplots showing protein preference based on cumulative licks in 5-min bins. Rows are individual rats. No bins are significantly different to 0.5 in nonrestricted rats (left) while all bins are significantly different in protein-restricted rats (right), † p < 0.05 vs. 0.5. (D) Cumulative licks for casein and maltodextrin. Protein-restricted rats show an increased intake of casein compared to maltodextrin from early in the session whereas no difference in intake is observed in nonrestricted rats. Lines are mean and shaded area is SEM.

Next, we analysed the temporal pattern of licking behaviour during the 1-hour session in each group of rats, to examine dynamics of casein and maltodextrin consumption (Fig. 1C and 1D). Visual inspection of cumulative licks over 60 minutes suggested that protein-restricted rats consumed more casein than maltodextrin throughout the entire session and that this difference began in the first few minutes of access to solutions. Conversely, non-restricted rats appeared to drink similar amounts of each solution. To quantify this difference, we examined if the slopes of cumulative casein and maltodextrin intake differed within each dietary group. We calculated the rate of change of licking (slope) for each rat for each solution and used two-way repeated measures ANOVA with Diet and Solution as factors. This analysis revealed a significant effect of Solution (F(1, 26) = 5.02, p = 0.034), no main effect of Diet (F(1,26 = 1.21, p = 0.281), but a significant Diet x Solution interaction (F(1,26) = 4.65, p = 0.040). Post hoc Sidak’s tests showed that the slopes for casein and maltodextrin differed for PR rats (p = 0.042) but not for NR rats (p = 1.00). To further assess the time course of casein preference, we compared protein preference in 5-minute bins throughout the session and found that preference was significantly above 0.5 in all bins for protein-restricted rats and non-restricted rats did not show a preference in any bins (one sample t-tests vs. 0.5, all ps < 0.05 with false discovery rate (FDR) correction). In summary, casein preference in protein-restricted rats arises quickly – within the first 5 minutes - after only minimal experience with nutrient-rich solutions.

### 3.2 Initial protein preference is not driven by elevated palatability of protein, but it is associated with an increase in protein’s palatability over the session

Next, we performed analysis of licking microstructure (Fig. 2), to assess whether the palatability of protein-containing solution was affected by protein restriction. As such, licks were divided into clusters with interlick intervals >500 ms. While the number of clusters initiated is thought to reflect post-ingestive feedback and satiety, the size of a cluster is considered an index of solution palatability [29,30].

**Figure 2.**
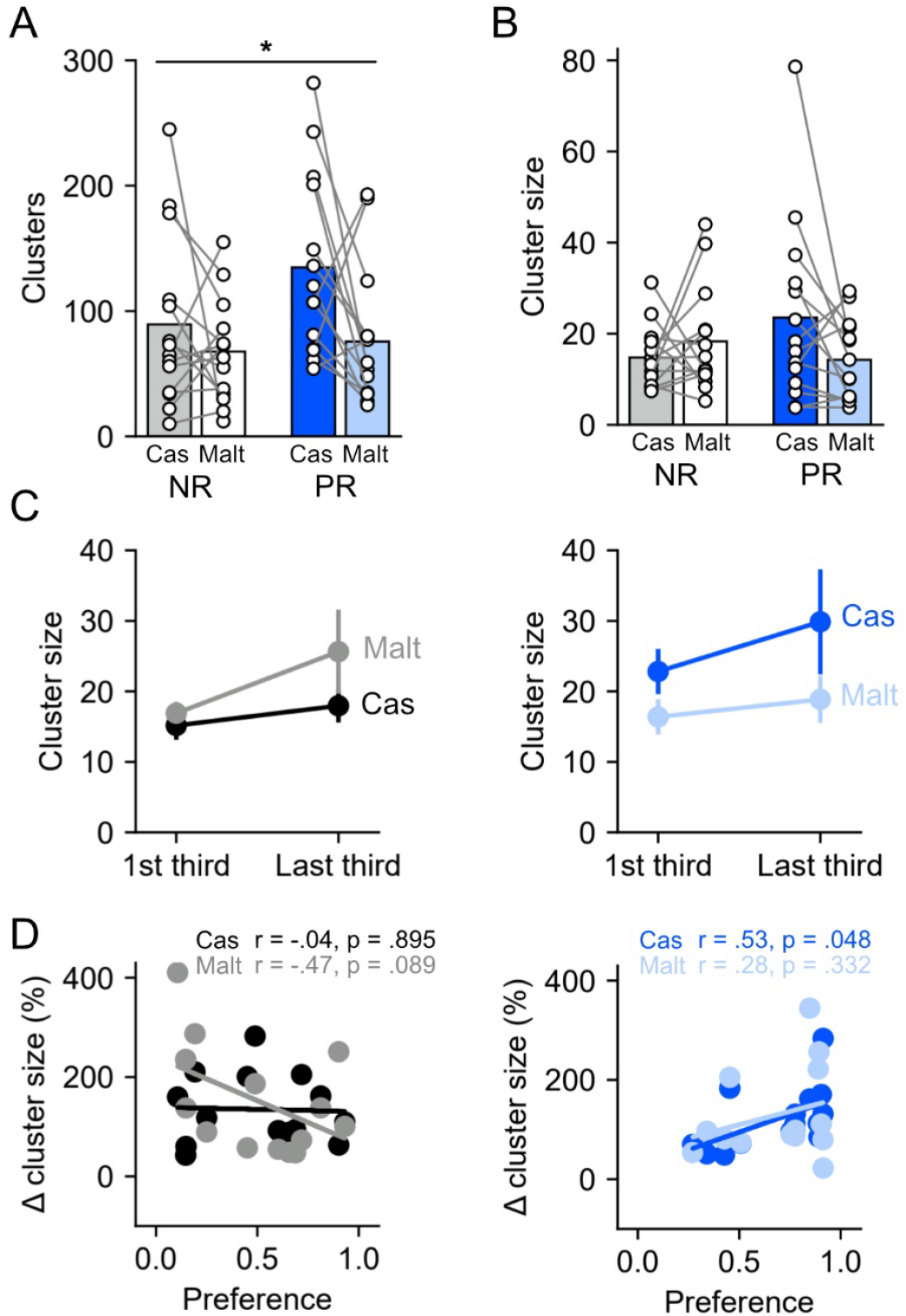
Payability of protein relative to carbohydrate during the first session is affected by protein restriction. (A) Number of clusters for casein and maltodextrin in non-restricted and protein-restricted rats. Bars are mean and circles are individual rats. *, p < 0.05 main effect of Solution. (B) Mean cluster size for casein and maltodextrin in nonrestricted and protein-restricted rats. Plotting conventions as in A. (C) Comparison of cluster sizes between the first third and the last third of the clusters for casein and maltodextrin in nonrestricted (left) and protein-restricted (right) rats. No significant difference is observed in either group for either solution, despite a tendency of greater cluster size for casein in protein-restricted rats (p = 0.073). Circles are mean and error bars show SEM. (D) Casein preference of non-restricted (left) and protein-restricted (right) rats plottedb as a function of percentage change in cluster sizes of casein and maltodextrin. In protein-restricted rats, casein preference positively correlates with change in cluster size of casein and there is a non-significant negative correlation between casein preference and change in cluster size for maltodextrin in non-restricted rats. Circles are individual rats and lines show the least squares fit.

We first analysed cluster number and size across the entire 1 h session in which rats experienced nutrient-containing solutions for the first time. Interestingly, cluster number was only somewhat consistent with our data on total licks (Fig. 2A). In fact, two-way ANOVA revealed a main effect of Solution (F(1, 26) = 6.12, p = 0.020), but no effect of Diet (F(1, 26) = 2.72, p = 0.111) and no significant Solution x Diet interaction (F(1, 26) = 1.31, p = 0.263). Subsequent *post hoc* comparisons suggested that the effect of Solution was predominantly driven by increased number of clusters for casein, relative to maltodextrin, in protein-restricted rats. However, this effect was weaker than that seen for total licks as it failed to reach significance after correction for multiple comparisons (uncorrected: p = 0.035; Sidak: p = 0.134). In non-restricted rats, there was no evidence for a difference between the solutions (uncorrected: p = 0.316; Sidak: p = 0.781).

We then analysed cluster size as a proxy for palatability of the solutions. When considering the entire 1 h session, we found that cluster size was not strongly affected by protein-restricted state in the first session when the nutrient-containing solutions were available – only the interaction term showed a trend towards significance (Fig. 2B; two-way repeated measures ANOVA: Solution (F(1, 26) = 0.78, p = 0.386), Diet (F(1, 26) = 0.41, p = 0.527), Solution x Diet interaction (F(1, 26) = 3.88, p = 0.059). We also considered whether the cluster size would change over time within the 1 h session, due to either positive post-ingestive effects or satiety and so divided bursts into thirds and compared average cluster size of the first vs. the last third [38] (Fig. 2C). Although visual inspection suggested that cluster size for casein increased during the session in protein-restricted rats while maltodextrin cluster size increased for non-restricted rats, a three-way repeated-measures ANOVA reported no significant main effects or interactions (all ps > 0.05).

To further investigate whether a positive change in the palatability of casein was associated with the preference for casein shown by protein-restricted rats, we examined whether, for individual rats, casein preference correlated with the change in cluster size from the first third to the last third of the clusters (Fig. 2D). In non-restricted rats, there was a weak negative correlation between preference for casein and the percentage change in cluster size for maltodextrin (r = -0.47, p = 0.089), while no correlation was observed for casein (r = -0.04, p = 0.895). These data indicate that, in non-restricted rats, a low preference for casein (i.e., a high preference for maltodextrin) was associated with an increase in the palatability of maltodextrin over the session. In contrast, in protein-restricted rats, casein preference was positively correlated with the change in cluster size for casein (r = 0.53, p = 0.048), while no correlation was observed for maltodextrin (r = 0.28, p = 0.332). Thus, in protein-restricted rats, the preference for casein shown during the session was associated with an increase in the palatability of casein over the session.

### 3.3 Protein preference in protein-restricted rats is persistent over consecutive test sessions and is associated with increased palatability of protein

After the first preference test session, four additional preference tests were conducted on consecutive days, to assess the evolution of casein preference over time (Fig. 3). A two-way repeated-measures ANOVA on casein preference revealed a main effect of Day (F(4, 104) = 3.69, p = 0.017), Diet (F(1, 26) = 15.49, p < 0.001) but no significant Day x Diet interaction (F(4, 104) = 0.45, p = 0.769), demonstrating a gradual change in casein preference in both groups (Fig 3A), but with protein-restricted rats exhibiting a greater casein preference throughout all tests compared to non-restricted rats. This heightened preference by protein-restricted rats was confirmed also by the one-sample t-tests performed on each Test Day (Test 2: t(13) = 9.35, p < 0.001; Test 3: t(13) = 10.24, p < 0.001; Test 4: t(13) = 2.08, p = 0.058; Test 5: t(13) = 8.41, p < 0.001). Conversely, non-restricted rats manifested a preference for casein only during the second preference test (Test 2: t(13) = 3.76, p = 0.002; Test 3: t(13) = 0.97, p = 0.348; Test 4: t(13) = 0.07, p = 0.943; Test 5: t(13) = 1.60, p = 0.135).

**Figure 3.**
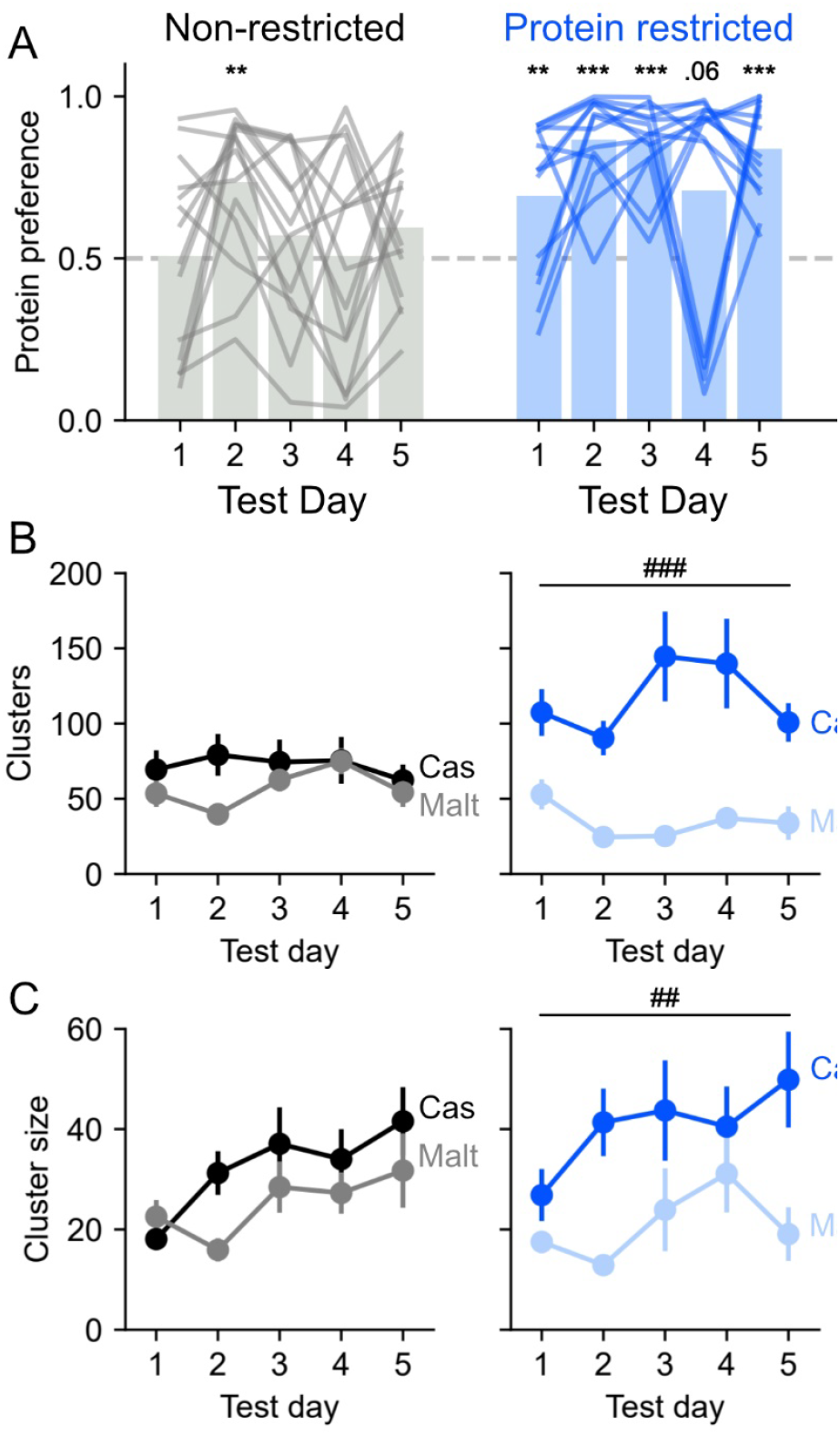
Protein preference in protein-restricted rats is persistent over consecutive test sessions and is associated with increased palatability of protein. After the first preference test, four consecutive preference tests were conducted. (A) Over the five preference tests, protein-restricted rats exhibit a greater casein preference compared to non-restricted rats, and this preference in protein-restricted rats is persists over repeated exposures to the solutions. (B) When all five sessions are considered, non-restricted rats (left) make the same number of clusters for casein and maltodextrin, while protein-restricted rats (right) make a greater number of clusters for casein compared to maltodextrin. (C) In non-restricted rats, the cluster size of both solutions moderately increased across the five test sessions; in protein-restricted rats, cluster size of casein was greater than maltodextrin and this was stable across the five sessions. Circles are mean and error bars show SEM. **, *** p < 0.01, 0.001 vs. 0.5 (one sample t-test). ##, ### p < 0.01, 0.001 (main effect of Solution).

As regards the number of clusters, three-way ANOVA revealed a main effect of Solution (F(1, 26) = 32.09, p < 0.001) and a significant Solution x Diet interaction (F(1, 26) = 15.17, p < 0.001). Follow-up, two-way repeated measures ANOVA on each diet group showed that non-restricted rats made similar number of clusters for casein and maltodextrin throughout the five test sessions (Fig. 3B; Day: F(4, 52) = 1.48, p = 0.221, Solution: F(1, 13) = 1.84, p = 0.198, Day x Solution F(4, 52) = 1.24, p = 0.305). In contrast, protein-restricted rats showed a greater number of clusters for casein compared to maltodextrin during all sessions (Solution: F(1, 13) = 39.72, p < 0.001, Day: F(4, 52) = 1.26, p = 0.299, Day x Solution interaction: F(4, 52) = 1.56, p = 0.199). Thus, when all five sessions were considered, consistent with their protein preference, the number of clusters for casein was greater than for maltodextrin in protein-restricted rats, but this did not change with repeated exposure.

We conducted a similar analysis but considering cluster size across all sessions. Three way ANOVA indicated a main effect of Day (F(2.6, 66.9) = 5.60, p = 0.003), of Solution (F(1, 26) = 14.33, p < 0.001) and a trend towards a Solution x Diet interaction (F(1, 26) = 3.05, p = 0.093). Follow-up two way repeated measures ANOVA showed that, in non-restricted rats, the size of lick clusters increased throughout the days of testing (Fig. 3C; Day: F(1.9, 24.7) = 4.14, p = 0.030, Solution: F(1, 13) = 2.49, p = 0.139, Day x Solution: F(4, 52) = 1.18, p = 0.332), indicating that the palatability of both solutions increased across the five test sessions. Interestingly, in protein-restricted rats, lick cluster size for both solutions remains relatively stable throughout the sessions, with cluster size for casein greater than for maltodextrin (Solution: F(1, 13) = 13.14, p = 0.003, Day: F(2.2, 28.1) = 2.13, p = 0.134, Day x Solution: F(2.3, 29.6) = 0.88, p = 0.436). This result indicates that palatability of casein was greater than maltodextrin in protein-restricted rats and this was relatively stable across the five sessions.

To summarize, these results suggest that palatability of casein is greater than maltodextrin in protein-restricted rats and this is stable across sessions. Conversely, non-restricted rats did not display an elevated palatability for either solution compared to the other, despite the palatability of both solutions moderately increasing over the course of testing.

### 3.5 Prior experience with casein-containing food does not influence initial protein preference (Experiment 2)

Finally, we wanted to confirm that the initial preference for casein that we observed on the first test day was not caused by previous exposure to casein through the low protein diet in protein-restricted, but not in non-restricted, rats. We tested this with a new cohort of rats in which the control non-restricted rats had extensive exposure to both the low protein diet and the control diet before their first preference test. Consistent with Experiment 1, we found that protein-restricted rats showed a significant preference for casein (one sample t-test vs. 0.5, t(7) = 4.35, p = 0.003) while non-restricted rats showed no preference (t(7) = 1.04, p = 0.332; Fig. 4A). Moreover, visual inspection of the cumulative licks throughout the session suggested a rapid increase in the number of licks for casein in protein-restricted rats only (Fig. 4B). These results suggest that familiarity with casein-containing food does not account for the rapid casein preference shown by protein-restricted rats.

**Figure 4.**
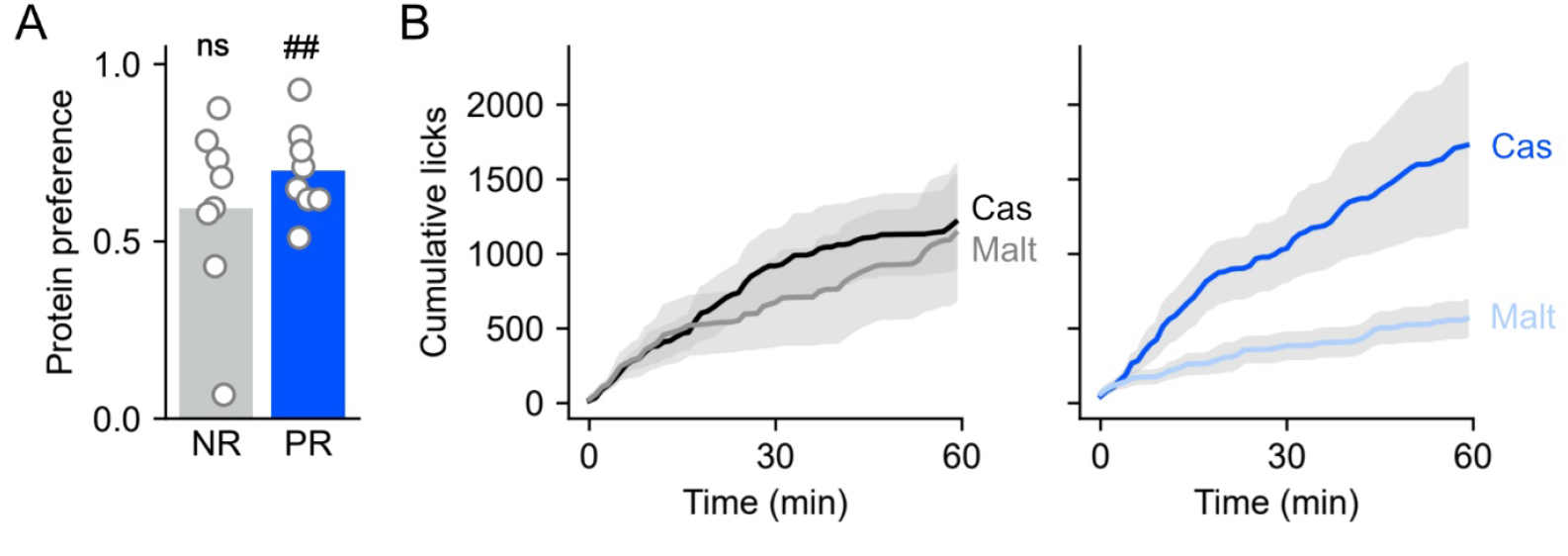
Previous experience with casein-containing food does not facilitate initial protein preference. Protein-restricted rats, as opposite to non-restricted, were maintained on a diet whose protein source came from casein. To assess whether familiarity with casein food fostered initial protein preference, a new batch of non-restricted rats was given the opportunity to consume casein-containing food before assessing initial casein preference. (A) Consistent with Experiment 1, protein-restricted rats showed a significant preference for casein, while non-restricted rats showed no preference. (B) Cumulative licks for casein and maltodextrin during the 60-minute preference test indicates an increased intake of casein compared to maltodextrin in protein-restricted rats only from the first minutes of exposure to solutions. ## p < 0.01 vs. 0.5 (one-sample t-test).

## 4 Discussion

In trying to understand the neural and cognitive mechanisms underlying protein appetite, an important unsolved question is whether the adaptive response of protein-seeking is innate or learning-dependent. Here, we assessed whether protein restriction affected the preference and palatability of a completely novel protein-containing solution, compared to novel carbohydrate. We found that, in response to protein need, protein preference arises rapidly in the face of a choice between protein vs. carbohydrate and is somewhat associated with an increase in protein palatability over the course of the first preference session, which is maintained over several days. In Experiment 2, we demonstrated that the rapid preference for protein-containing solution (casein) is not just driven by the familiarity with casein-containing food but instead is dependent on the state of protein restriction.

Our study supports evidence from our previous observations [10,15] that protein-restricted rats develop a robust preference for protein-containing solution (casein) over carbohydrate (maltodextrin) when given the choice between the two macronutrients. However, in our previous experiments, rats experienced nutrient-containing solutions in a block of conditioning sessions before the preference test, allowing flavour-nutrient associations to be learned via post-ingestive signalling in a specific physiological state (non-restricted or protein-restricted) [39]. This paradigm prevented us from discriminating whether protein drive was innate or involved some form of learning. Thus, in the present study, the conditioning block was removed, and initial protein preference was assessed after ∼10 days of protein restriction without rats first experiencing the test solutions. Despite these changes, protein-restricted rats still manifested a strong casein preference apparent in the very first session. Interestingly, if protein intake is examined in situations when only a single solution is available at the time, rather than with two choices, effects of protein restriction are often not seen. As such, in our previous study when only one type of solution was available during conditioning, there was no difference in consumption between the first day of casein and the first day of maltodextrin [10,13,15]. Moreover, when the preference test consisted of both forced-choice trials (one solution at a time, casein or maltodextrin) and free-choice trials (casein vs. maltodextrin) within the same session, protein-restricted animals manifested a preference for casein during free-choice trials only [10,13,15]. It is therefore possible that the casein preference manifested in the present study is due to a negative contrast effect, leading to a decreased value of maltodextrin, relative to casein, as a function of the comparison between the two solutions [40].

Analysis of cumulative licks over the session showed that casein preference in protein-restricted rats arose within the first 5 minutes of the preference test. The speed of this effect is consistent with one previous study showing that, during protein need, rats took just one minute to manifest a preference for protein (i.e., fibrin and ovalbumin) over carbohydrate [21]. The best-documented example of a rapid nutrient-specific appetite is the immediate onset of sodium appetite that arises under sodium depletion [41], which has been interpreted as the result of innate mechanisms making rats especially responsive to sodium taste during deficiency [42]. It is unlikely that rats had ever experienced protein restriction before the start of the experiment, therefore there were no previous opportunities to learn that consuming protein-rich food would alleviate protein deficiency. However, others have reported that lysine-deficient rats develop a lysine preference within 30 minutes, a time window that has been interpreted as the results of post-ingestive feedback [43]. Moreover, rapid post-oral effects (within 15 minutes) that increase glucose and fat ingestion have been reported in mice [44]. Future studies are therefore needed to explain the mechanisms allowing the rapid onset of protein preference during protein restriction. Experiments involving intra-gastric infusions of protein in different parts of the gastric tract and/or using sham-feeding procedures to remove post-ingestive signals while maintaining sensory signaling will need to be undertaken to disentangle the contribution of oral vs. post-oral mechanisms in protein appetite.

Our group has reported that the conditioned protein preference developed by protein restricted animals is associated with an increase in the palatability of casein-containing solution, based on the increased cluster size for casein, relative to maltodextrin [10,13,29]. In the present study, protein-restricted rats initiated more lick clusters for casein, without a significant increase in palatability during the first preference test. This is consistent with previous studies reporting that lysine-deficient rats increase the number of clusters for lysine-containing solution, but not cluster size [43]. Considering that the number of clusters can be considered an index of motivation to ingest a solution and is sensitive to post-ingestive signals [29], the present data are also consistent with our previous findings on the response of the dopamine system to protein appetite [16–18]. It is possible that the increased casein palatability shown by protein-restricted rats in our previous studies depends on repeated association, through conditioning, of the sensory characteristics of casein solution with the post-ingestive feedback of protein repletion. This possibility is strengthened by our observations that cluster size for casein was greater than for maltodextrin when all five consecutive daily sessions were considered. Reduction of palatability due to satiety within a single session was previously demonstrated, using either mimetic responses [45,46] or licking microstructure analysis [47]. During the first preference tests, the comparison of cluster size at the beginning and at the end of the session did not highlight any significant difference. However, when we considered individual performances, we found a positive correlation between casein preference and increased palatability of casein across the session in protein-restricted rats. These results support the idea that increased palatability of casein is not wholly responsible for expression of protein preference but may contribute and enhance the preference for it.

The low protein diet we used contained casein as its protein source whereas the control diet did not. Thus, in Experiment 1 the protein-restricted rats were not completely naïve to casein, raising the possibility that they consumed more casein than maltodextrin due to a decrease in novelty-induced neophobia. Using pre-exposure to casein-containing low protein diet, we showed that casein preference remained low in non-restricted animals compared to protein-restricted rats. This result was observed despite robust intake of the low protein diet during the pre-exposure phase, ruling out a potential role of neophobia for casein.

Here, we only gave rats the choice of casein as a protein source so whether protein-restricted rats would show rapid preference for other proteins is unclear. However, early findings showed that several proteins including gluten, ovalbumin, and fibrin could stimulate a protein preference in protein-restricted rats (interestingly, casein did not) [21,22]. In addition, a recent study showed equally strong protein preference for whey protein as casein [18], although here rapid effects of the protein were not tested. Understanding differences between proteins, especially those from different sources (e.g., animal-derived vs. plant-derived), will be useful for determining implications for human health.

When rodents are sodium depleted, changes in sodium gustatory perception have been reported at the level of taste receptors on the tongue and palate epithelium [48] and the chorda tympani nerve [49], and alter responses to sodium taste. Biological modifications in the periphery may explain the immediate responsiveness of sodium-deprived animals to sodium-containing food. Protein restriction has been shown to cause physiological changes in taste and olfaction, but the underlying mechanisms remain unclear. One possibility is that modifications in signalling pathways in the gustatory system, or upregulation of taste receptors, might be responsible for increased protein sensing during protein restriction. For example, early dietary protein restriction in rats leads to alterations in the chorda tympani nerve, both in terms of responses to gustatory stimuli and in the volume of its terminal field [50]. Amino acids are known to be detected by specific receptors such as Tas1r2/Tas1r3 and mGluR4 [51,52], whose modifications might be a candidate mechanism. Olfactory function during protein restriction might be altered by modifications at the level of olfactory receptors, or even centrally (i.e., olfactory bulb). As such, protein-restricted mice showed oxidative stress in the olfactory epithelium and reduced mature olfactory sensory neurons [53]. Whether such changes are the results of post-ingestive feedback is still unknown and future investigations will be important to determine the underlying mechanisms.

In summary, here we show that protein-restricted rats rapidly show a preference for a flavoured protein solution, relative to a carbohydrate solution, even upon their first experience of it. This increased intake was not strongly associated with a change in motivation or satiety mechanisms, although there were indications that casein palatability increased in protein-restricted rats over time. Future studies will allow the mechanisms of this rapid preference to be further interrogated.

## Acknowledgements and funding

The authors acknowledge the help and support from the staff of the Division of Biomedical Services, Preclinical Research Facility, University of Leicester, for technical support and the care of experimental animals as well as colleagues in the Department of Neuroscience, Psychology and Behaviour at the University of Leicester for their academic contribution. This work was funded by the Biotechnology and Biological Sciences Research Council [grant #BB/M007391/1 to J.E.M.], the European Commission [grant #GA 631404 to J.E.M.], The Leverhulme Trust [grant #RPG-2017-417 to J.E.M. and J.A-S.], and Tromsø Research Foundation [grant #19-SG-JMcC to J.E.M.).

## References

[1] C.P. Richter, Total self regulatory functions in animals and human beings, Harvey Lect 38 (1943) 63–103.

[2] S.J. Simpson, D. Raubenheimer, The nature of nutrition: a unifying framework from animal adaptation to human obesity, Princeton University Press, Princeton, 2012.

[3] D. Mayntz, D. Raubenheimer, M. Salomon, S. Toft, S.J. Simpson, Nutrient-Specific Foraging in Invertebrate Predators, Science 307 (2005) 111–113. 10.1126/science.1105493.

[4] S.J. Simpson, D. Raubenheimer, Obesity: the protein leverage hypothesis, Obesity Rev 6 (2005) 133–142. 10.1111/j.1467-789X.2005.00178.x.

[5] B. Musten, D. Peace, G.H. Anderson, Food intake regulation in the weanling rat: self-selection of protein and energy, J Nutr 104 (1974) 563–572. 10.1093/jn/104.5.563.

[6] P.D. Leathwood, D.V. Ashley, Strategies of protein selection by weanling and adult rats, Appetite 4 (1983) 97–112. 10.1016/s0195-6663(83)80006-3.

[7] C.P. Richter, B. Barelare, Nutritional requirements of pregnant and lactating rats studied by the selfselection method, Endocrinology 23 (1938) 15–24. 10.1210/endo-23-1-15.

[8] A.I. Leshner, H.I. Siegel, G. Collier, Dietary self-selection by pregnant and lactating rats, Physiol Behav 8 (1972) 151–154. 10.1016/0031-9384(72)90144-8.

[9] T.J. Roberts, M.J. Azain, B.D. White, R.J. Martin, Rats treated with somatotropin select diets higher in protein, J Nutr 125 (1995) 2669–2678. 10.1093/jn/125.10.2669.

[10] M. Murphy, K.Z. Peters, B.S. Denton, K.A. Lee, H. Chadchankar, J.E. McCutcheon, Restriction of dietary protein leads to conditioned protein preference and elevated palatability of protein-containing food in rats, Physiol Behav 184 (2018) 235–241. 10.1016/j.physbeh.2017.12.011.

[11] E.L. Gibson, D.A. Booth, Acquired protein appetite in rats: Dependence on a protein-specific need state, Experientia 42 (1986) 1003–1004. 10.1007/BF01940706.

[12] C.M. Hill, T. Laeger, M. Dehner, D.C. Albarado, B. Clarke, D. Wanders, S.J. Burke, J.J. Collier, E. Qualls-Creekmore, S.M. Solon-Biet, S.J. Simpson, H.-R. Berthoud, H. Münzberg, C.D. Morrison, FGF21 Signals Protein Status to the Brain and Adaptively Regulates Food Choice and Metabolism, Cell Rep 27 (2019) 2934-2947.e3. 10.1016/j.celrep.2019.05.022.

[13] K.L. Volcko, J.E. McCutcheon, Protein preference and elevated plasma FGF21 induced by dietary protein restriction is similar in both male and female mice, Physiol Behav 257 (2022) 113994. 10.1016/j.physbeh.2022.113994.

[14] K.L. Volcko, H. Taghipourbibalan, J.E. McCutcheon, Intermittent protein restriction elevates food intake and plasma ghrelin in male mice, Appetite 203 (2024) 107671. 10.1016/j.appet.2024.107671.

[15] G. Chiacchierini, F. Naneix, K.Z. Peters, J. Apergis-Schoute, E.M.S. Snoeren, J.E. McCutcheon, Protein Appetite Drives Macronutrient-Related Differences in Ventral Tegmental Area Neural Activity, J Neurosci 41 (2021) 5080–5092. 10.1523/JNEUROSCI.3082-20.2021.

[16] G. Chiacchierini, F. Naneix, J. Apergis-Schoute, J.E. McCutcheon, Restriction of dietary protein in rats increases progressive-ratio motivation for protein, Physiol Behav 254 (2022) 113877. 10.1016/j.physbeh.2022.113877.

[17] C.-T. Wu, D. Gonzalez Magaña, J. Roshgadol, L. Tian, K.K. Ryan, Dietary protein restriction diminishes sucrose reward and reduces sucrose-evoked mesolimbic dopamine signaling in mice, Appetite 203 (2024) 107673. 10.1016/j.appet.2024.107673.

[18] M.S.H. Khan, S.Q. Kim, R.C. Ross, F. Corpodean, R.A. Spann, D.A. Albarado, S.O. Fernandez-Kim, B. Clarke, H.-R. Berthoud, H. Münzberg, D.H. McDougal, Y. He, S. Yu, V.L. Albaugh, P.L. Soto, C.D. Morrison, FGF21 acts in the brain to drive macronutrient-specific changes in behavioral motivation and brain reward signaling, Mol Metab 91 (2025) 102068. 10.1016/j.molmet.2024.102068.

[19] F. Naneix, K.Z. Peters, A.M.J. Young, J.E. McCutcheon, Age-dependent effects of protein restriction on dopamine release, NPP 46 (2021) 394–403. 10.1038/s41386-020-0783-z.

[20] B.G. Galef Jr., Is there a specific appetite for protein, in: Neural and Metabolic Control of Macronutrient Intake, CRC Press, 2000: pp. 19–26.

[21] J.A. Deutsch, B.O. Moore, S.C. Heinrichs, Unlearned specific appetite for protein, Physiology & Behavior 46 (1989) 619–624. 10.1016/0031-9384(89)90341-7.

[22] S.C. Heinrichs, J.A. Deutsch, B.O. Moore, Olfactory self-selection of protein-containing foods, Physiology & Behavior 47 (1990) 409–413. 10.1016/0031-9384(90)90101-9.

[23] V. Augustine, S. Lee, Y. Oka, Neural Control and Modulation of Thirst, Sodium Appetite, and Hunger, Cell 180 (2020) 25–32. 10.1016/j.cell.2019.11.040.

[24] D. Denton, The hunger for salt: an anthropological, physiological and medical analysis, 2. print. of 1. ed, Springer, Berlin Heidelberg, 1984.

[25] K.C. Berridge, F.W. Flynn, J. Schulkin, H.J. Grill, Sodium Depletion Enhances Salt Palatability in Rats, (n.d.).

[26] D. DiBattista, Effects of time-restricted access to protein and to carbohydrate in adult mice and rats, Physiology & Behavior 49 (1991) 263–269. 10.1016/0031-9384(91)90042-M.

[27] B.D. White, M.H. Porter, R.J. Martin, Protein selection, food intake, and body composition in response to the amount of dietary protein, Physiol Behav 69 (2000) 383–389. 10.1016/s0031-9384(99)00232-2.

[28] C.D. Morrison, T. Laeger, Protein-dependent regulation of feeding and metabolism, Trends in Endocrinology & Metabolism 26 (2015) 256–262. 10.1016/j.tem.2015.02.008.

[29] F. Naneix, K.Z. Peters, J.E. McCutcheon, Investigating the Effect of Physiological Need States on Palatability and Motivation Using Microstructural Analysis of Licking, Neuroscience 447 (2020) 155–166. 10.1016/j.neuroscience.2019.10.036.

[30] J.D. Davis, G.P. Smith, Analysis of the microstructure of the rhythmic tongue movements of rats ingesting maltose and sucrose solutions, Behav Neurosci 106 (1992) 217–228. 10.1037/0735-7044.106.1.217.

[31] Y. Benjamini, A.M. Krieger, D. Yekutieli, Adaptive linear step-up procedures that control the false discovery rate, Biometrika 93 (2006) 491–507. 10.1093/biomet/93.3.491.

[32] D.A. Fitts, Minimizing Animal Numbers: The Variable-Criteria Sequential Stopping Rule, Comparative Medicine 61 (2011).

[33] D.A. Fitts, Improved stopping rules for the design of efficient small-sample experiments in biomedical and biobehavioral research, Behavior Research Methods 42 (2010) 3–22. 10.3758/BRM.42.1.3.

[34] W. McKinney, Data Structures for Statistical Computing in Python, in: Austin, Texas, 2010: pp. 56–61. 10.25080/Majora-92bf1922-00a.

[35] J.D. Hunter, Matplotlib: a 2D graphics environment, Computing in Science & Engineering 9 (2007) 90–95. 10.1109/MCSE.2007.55.

[36] R. Vallat, Pingouin: statistics in Python, JOSS 3 (2018) 1026. 10.21105/joss.01026.

[37] M. Waskom, seaborn: statistical data visualization, JOSS 6 (2021) 3021. 10.21105/joss.03021.

[38] A.C. Spector, S.J. St. John, Role of taste in the microstructure of quinine ingestion by rats, American Journal of Physiology-Regulatory, Integrative and Comparative Physiology 274 (1998) R1687–R1703. 10.1152/ajpregu.1998.274.6.R1687.

[39] A. Sclafani, Post-ingestive positive controls of ingestive behavior, Appetite 36 (2001) 79–83. 10.1006/appe.2000.0370.

[40] C. Mitchell, C. Flaherty, Temporal dynamics of corticosterone elevation in successive negative contrast, Physiology & Behavior 64 (1998) 287–292. 10.1016/S0031-9384(98)00072-9.

[41] P.J. Handal, Immediate acceptance of sodium salts by sodium deficient rats, Psychon Sci 3 (1965) 315–316. 10.3758/BF03343156.

[42] G. Wolf, Innate mechanisms for regulation of sodium intake., in: Olfaction and Taste, Rockefeller University Press, 1969: pp. 548–553.

[43] S. Markison, A.C. Spector, B.L. Thompson, J.C. Smith, Time Course and Pattern of Compensatory Ingestive Behavioral Adjustments to Lysine Deficiency in Rats, The Journal of Nutrition 130 (2000) 1320–1328. 10.1093/jn/130.5.1320.

[44] S. Zukerman, K. Ackroff, A. Sclafani, Rapid post-oral stimulation of intake and flavor conditioning by glucose and fat in the mouse, AJP-RICP 301 (2011) R1635–R1647. 10.1152/ajpregu.00425.2011.

[45] K.C. Berridge, Modulation of taste affect by hunger, caloric satiety, and sensory-specific satiety in the rat, Appetite 16 (1991) 103–120. 10.1016/0195-6663(91)90036-R.

[46] L.A. Eckel, K.P. Ossenkopp, Cholecystokinin reduces sucrose palatability in rats: evidence in support of a satiety effect, Am J Physiol 267 (1994) R1496–1502. 10.1152/ajpregu.1994.267.6.R1496.

[47] J.D. Davis, G.P. Smith, B. Singh, A microstructural analysis of the control of water and isotonic saline ingestion by postingestional stimulation, Physiol Behav 66 (1999) 543–548. 10.1016/s0031-9384(98)00325-4.

[48] J. Chandrashekar, C. Kuhn, Y. Oka, D.A. Yarmolinsky, E. Hummler, N.J.P. Ryba, C.S. Zuker, The cells and peripheral representation of sodium taste in mice, Nature 464 (2010) 297–301. 10.1038/nature08783.

[49] R.J. Contreras, M. Frank, Sodium deprivation alters neural responses to gustatory stimuli., The Journal of General Physiology 73 (1979) 569–594. 10.1085/jgp.73.5.569.

[50] J.E. Thomas, D.L. Hill, The effects of dietary protein restriction on chorda tympani nerve taste responses and terminal field organization, Neuroscience 157 (2008) 329–339. 10.1016/j.neuroscience.2008.09.013.

[51] E.R. Liman, Y.V. Zhang, C. Montell, Peripheral Coding of Taste, Neuron 81 (2014) 984–1000. 10.1016/j.neuron.2014.02.022.

[52] N. Chaudhari, E. Pereira, S.D. Roper, Taste receptors for umami: the case for multiple receptors, The American Journal of Clinical Nutrition 90 (2009) 738S–742S. 10.3945/ajcn.2009.27462H.

[53] A. Tuerdi, S. Kikuta, M. Kinoshita, T. Kamogashira, K. Kondo, T. Yamasoba, Zone-specific damage of the olfactory epithelium under protein restriction, Sci Rep 10 (2020) 22175. 10.1038/s41598-020-79249-3.

