## Supplemental Figures for "Dietary protein restriction in rats leads to a rapid within-session preference for protein"

**Supplemental Figures for Chiaccherini et al.**

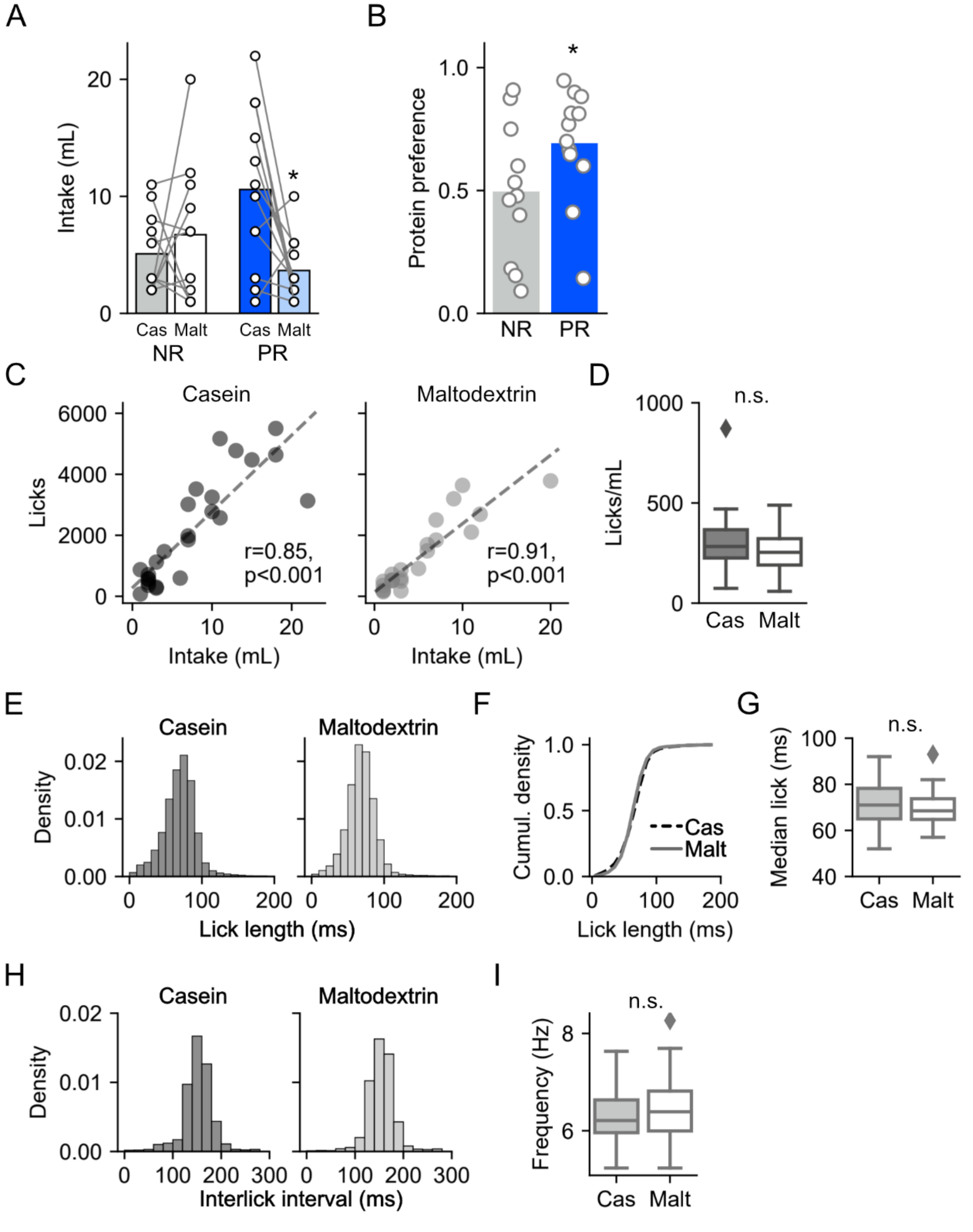


**Supplemental Figure 1. Licking behaviour is a good proxy for intake.** (A) Preference for casein over maltodextrin is still seen in protein-restricted rats when intake is measured by weighing bottles before and after each session instead of using licks. Bars are mean and circles are individual rats. *, $p < 0.05$ vs. casein Sidak post-hoc after significant 2-way interaction ($F_{1,21} = 7.31$, $p = 0.013$). (B) Similarly, when protein preference is calculated using intake instead of licks, a preference is still seen in protein-restricted rats but not in non-restricted rats (PR: $t_{11} = 2.91$, $p = 0.014$; $t_{10} = 0.07$, $p = 0.944$). (C) A strong correlation is seen between licks and intake for both casein and maltodextrin solutions. (D) No difference in licks per mL intake was seen between casein and maltodextrin solutions ($t_{46}$ = 1.03, p = 0.309). Boxplots show distribution across all rats. (E, F) Histograms show distribution of lick lengths for each solution and cumulative probability shows that >98% of licks are <200 ms demonstrating that contact lickometers were measuring individual licks with high fidelity. (G) There was no difference in median lick length between casein and maltodextrin ($t_{26}$ = 0.66, p = 0.514). (H) Histograms show interlick intervals within bursts. (I) No difference between intraburst frequency when licking for casein or maltodextrin ($t_{26}$ = 1.49, p = 0.149).

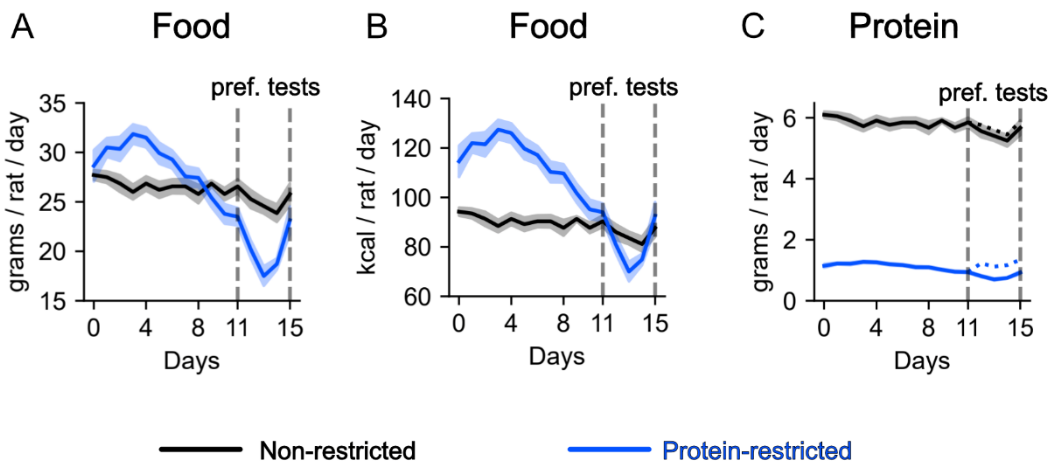


**Supplementary Figure 2. Approximate food intake during the experiment.** (A) Food intake in grams. Dashed lines show time period during which daily preference test sessions took place. Note that as rats were group-housed, food intake is approximated by dividing by number of rats in cage. (B) Food intake in kcals. Plotting conventions as in A. (C) Protein intake was approximated from food intake. In addition, the dotted lines that begin on day 11 when preference tests start show protein intake combined between test sessions and home cage intake. Solid lines are mean and shaded area is SEM.
